# Variant pathogenic prediction by locus variability, the importance of the last picture of evolution

**DOI:** 10.1101/2020.11.06.371195

**Authors:** Jose Luis Cabrera-Alarcon, Jose Antonio Enriquez, Jorge Garcia Martinez, Fátima Sánchez-Cabo

## Abstract

Accurate pathogenic detection for single nucleotide variants (SNVs) is a key problem to perform variant ranking in whole exome sequencing studies. Several in silico tools have been developed to identify deleterious variants. Locus variability, computed as Shannon entropy from gnomAD/helixMTdb variant allele frequencies can be used as pathogenic variants predictor. In this study we evaluate the use of Shannon entropy in non-coding mitochondrial DNA and also in coding regions with an additional selective pressure other than that imposed by the genetic code, as are splice-sites. To benchmark this functionality in non-coding mitochondrial variants, Shannon entropy was compared with HmtVar disease score, outperforming it in non-coding SNVs (AUC_H_=0.99 in ROC curve and PR-AUC_H_=1.00 in Precision-recall curve). In the same way, for splice-sites’ variants, Shannon entropy was compared against two state-of-the-art ensemble predictors ada score and rf score, matching their overall performance both in ROC curves (AUC_H_=0.95) and Precision-recall curves (PR-AUC=0.97). Therefore, locus variability could aid in variant ranking process for these specific types of SNVs.

## BACKGROUND

Whole Exome Sequencing (WES) is a powerful technique used in the frame of genetic based diseases, specially for mendelian diseases. However, given the high degree of variability in human population, WES renders a large amount of variants, making a challenge discerning pathogenic variants from the vast neutral background. For this purpose, researchers have built several predictors to aid in variant ranking process, for the detection of deleterious variants. In this sense, the degree of human variability could be used as pathogenic predictor.

Up to date the greatest effort in gathering human variability is presented in gnomAD, where are included variants detected in 125748 whole exome sequences and 15708 whole genome sequences from unrelated individuals^1^. Similarly, helixMTdb is a database that retrieve human genetic variability in mitochondrial DNA from 196,554 unrelated individuals^2^. All this information could be integrated as a measurement of locus-wise variability, that gives an estimation about the mutational freedom by genomic position. Furthermore, this measurement could reflect directly the relative importance of a specific genomic position and therefore the pathogenic status of any mutation placed there. Nevertheless, such predictor will define a genomic position more than a specific mutation. It is known that a characteristic of the genetic code that vast majority of amino acids can be translated by more than one codon. This redundancy results in the coexistence at the same genomic positions of synonymous and lethal SNVs, translated as a decrease in the accuracy of Shannon entropy as predictor.

On the other hand, this kind of predictor could be useful for genomic positions where there is no influence of the genomic code, as non-coding regions, or where the effect of genetic code redundancy is eclipsed by the selective pressure imposed by an additional functionality, as the splicing in splice-sites.

Eukaryotic cells harbor two different genomes, the nuclear and the mitochondrial DNA. Both genomes have their own evolutionary engines: while nuclear genome presents sexual reproduction as source of variability with sister chromatid exchange, mitochondrial DNA is mainly maternally inherited and has a higher mutation rate as its main source of variability. Therefore, mitochondrial DNA has its own conservation path and population frequencies that may not present the same behavior as nuclear DNA for these features. In this context, some of the variants produced by this larger mutational rate, may lead to mitochondrial disorders. Hence, in variants deleteriousness detection, this mitochondrial particularities are a major point to take into account, not only for protein-coding variants, where are focused the vast majority of developed predictors, but also for non-coding sites as Control region, t-RNA genes or r-RNA genes, that represent approximately the 32% of this genome.

At the same time, alternative splicing (AS) is a major biological mechanism for rising protein diversity in organisms. So it should come at no surprise that complexity in AS is correlated with evolution and tissue complexity^3^. Given the biological nature of splicing process, splice regions, understood as those positions located at the boundary of an exon and an intron (splice-site), had a different evolutionary process than non-splice-sites. Therefore, these regions may not have the same freedom to be mutated than other protein-coding sites, situation that can be translated in the degree of variability observed in human population.

Here, we propose the use of Shannon entropy calculated over the population frequencies associated to a genomic position, to detect pathogenic single nucleotide variants (SNVs) placed in splice-sites and non-coding regions in mitochondrial DNA.

## MATERIAL AND METHODS

### Datasets

In this study we used a composed dataset, that contains 131043 unique variants (65775 pathogenic, 65268 neutral). These variants were obtained from five independent benchmark datasets *HumVar^4^, ExoVar^5^, VariBench^6^, predictSNP^7^ and SwissVar^8^* and also variants selected from Clinvar archive^9,10^, classified as benign or pathogenic variants. These 131043 were split according to splice-site/not-splice-site location, considering splice-sites either within 1-3 bases of the exon or 3-8 bases of the intron, resulting in 120744 not-splice-site variants and 10294 splice-site SNVs. In order to benchmark the use of Shannon entropy for deleteriousness detection in splice-site SNVs, a subset of 7941 out of 10294 variants, for which there were pre-computed values for ada score and rf score in dbNSFP^11^, was selected.

For mitochondrial evaluation of Shannon entropy, mitochondrially encoded variants were selected from the above datasets joined with confirmed variants described in Mitomap^12^ as pathogenic plus high frequency variants (Fq>0,01) described in this database. A total number of 451 variants were obtained, 169 non-coding SNVs (80 in tRNA genes, 33 in rRNA genes and 28 in control region) and 282 protein-coding SNVs. In non-coding mitochondrial variants 64 out of 169 were neutral and 105 were pathogenic. In addition, in protein-coding mitochondrial variants 157 were neutral and 125 were pathogenic. These variants were annotated with HmtVar disease score^13^, retrieving 336 annotated variants (112 non-coding & 224 protein-coding), for benchmarking Shannon entropy.

### Variants annotation

Splice-site condition was annotated using Variant effect Predictor^14^, which also was used to retrieve ada score and rf score values from dbNSFP. Beside this, phastCons^15^ and phylop^16^ conservation scores calculated on multiple sequence alignment from sequences of 100 species of vertebrates, was retrieved using UCSC table browser data retrieval tool^17^. Variant allele frequencies were retrieved from gnomAD 2.1.1 for nuclear encoded variants or helixMTdb, for mitochondrial SNVs. Finally, HmtVar disease scores were obtained using HmtVar API.

### Shannon entropy

Locus variability was calculated using gnomAD/helixMTdb variant allele frequencies as Shannon entropy for the genomic positions of variants considered in our data sets following the expression:

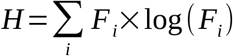

where H is Shannon entropy and F_i_ is variant allele frequency observed in gnomAD or helixMTdb for each allele observed in a specific genomic position. This parameter can take values from 0 onwards, meaning the higher the value the more variability observed for that genomic position where is placed the variant we are studying.

### Benchmark of Shannon entropy predictors for splice-site SNVs

Shannon entropy performance was compared in this variants with two state-of-the-art ensemble scores, **ada score** and **rf score** and also with two conservation scores as **phyloP** and **phastCons**, based on AUC/PR-AUC in receiving operating characteristic (ROC) curves and precision recall (PR) curves. Joined with this, non parametric statistical hypothesis contrasts were performed to test AUC differences for ROC curves, calculating the D-statistic as:

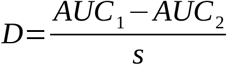

where AUC_1_ is the area under the curve of one predictor, AUC_2_ is the area under the curve of the other predictor and **s** is the standard deviation for the difference between both values in 2000 subsets obtained by bootstrap re-sampling, as described in pROC R-package^18^, which was used for this purpose. ROC and PR curves analysis were performed using ROCR^19^ and precrec^20^ R-packages.

### Benchmark of Shannon entropy in mitochondrial SNVs

Shannon entropy was compared against HmtVar disease score for pathogenic prediction in non-coding SNVs. In addition, Shannon entropy will also be evaluated in protein-coding

## RESULTS

It was observed a significant difference for Shannon entropy between variants located in splice and not splice regions, by non parametric Kolmogorov-Smirnov test (D=0.186, p<0.001), figure 1a.

**Figure 1.**
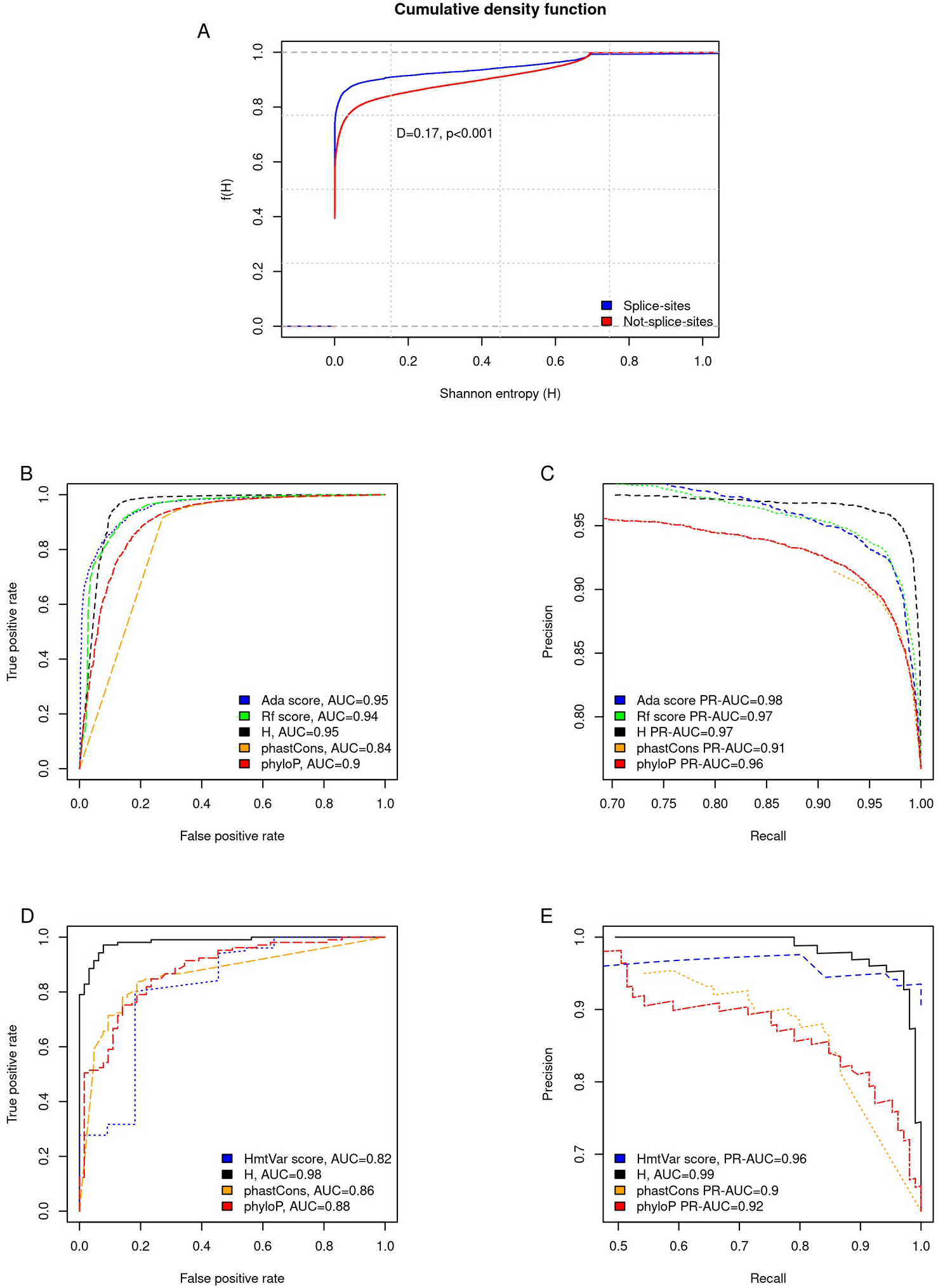
Cumulative density function of locus variability as Shannon entropy comparing splice-site and not-splice-site positions (A). ROC curves (B) and PR curves (C) for Shannon entropy, ada score, rf score, phastCons and phyloP conservation scores for deleteriousness detection in splice-sites SNVs. ROC curves (D) and precision-recall curves (E) for Shannon entropy, HmtVar disease score phastCons and phyloP conservation scores for pathogenic variants detection in mitochondrial non-coding SNVs. D-statistic and its associated p-value refer to Kolmogorov-Smirnov non parametric test. Abreviations: receiving operating characteristic (ROC), precision-recall (PR), Shannon entropy (H) and single nucleotide variants (SNVs).

Regarding to the benchmark results in splice-sites, Shannon entropy, with an AUC_H_=0.95, outperformed rf score (AUC_rf_score_=0.94), phastCons (AUC_phastCons_= 0.84) and phyloP (AUC_phyloP_= 0.9) and matches the results of ada score (AUC_ada_score_=0.95), figure 1b. AUC differences between Shannon entropy and rf score, phastCons or phyloP were statistically significant (D_H_rf_score_=2.02, p<0.05; D_H_phastCons_=19.12, p<0.001 and D_H_phyloP_=9.10, p<0.001), while there were no difference with ada score (D_H_ada_score_=-0.35, p=0.7232). Regarding to PR curves outcomes, Shannon entropy presented a PR-AUC_H_=0.97 outperforming to phastCons (PR-AUC_phastCons_=0.91) and phyloP (PR-AUC_phyloPs_=0.96), matching rf score results (PR-AUC_rf_score_=0.97) and being slightly surpassed by ada score (PR-AUC_rf_score_=0.98), figure 1c.

On the other hand, overall performance of Shannon entropy in not-splice-site SNVs, rendered an AUC=0.86.

For mitochondrial non-coding SNVs, Shannon entropy presented the best behavior compared with HmtVar score, phastCons or phyloP, both in ROC curve (AUC_H_=0.98, AUC_HmtVar_=0.82, AUC_phastCons_=0.86 & AUC_phyloP_=0.88) and PR curve (PR-AUC_H_=1.00, PR-AUC_HmtVar_=0.99, PR-AUC_phastCons_=0.9 & PR-AUC_phyloP_=0.92), figures 1d and 1e. These differences between Shannon entropy and all other score in non-coding variants, were statistically significant (D_H_HmtVar_=2.34, p<0.05; D_H_phastCons_=4.31, p<0.001 & D_H_phyloP_=3.99, p<0.001).

To the contrary, Shannon entropy presented a poor performance in protein coding variants, AUC_H_=0.68, surpassed by HmtVar disease score (AUC_HmtVar_=0.78).

According to our data, based on Youden index, we propose a cutoff for deleteriousness detection in splice variants for Shannon entropy H≤0.001053301 (sensitivity=0.97, specificity=0.87), values below this cutoff should be considered pathogenic. On the other hand, for mitochondrial non-coding SNVs we propose a threshold H≤0.009523692 (sensitivity=0.97, specificity=0.92).

## DISCUSSION

Shannon entropy, computed over gnomAD or helixMTdb variant allele frequencies, reflects the freedom for a specific genomic position to be mutated in the population, showing the final picture of selective pressures. According to our outcomes, locus variability in splice-site positions is significantly lower than in not splice-site positions, confirming the assumption of a different performance between both types of genomic regions. This different behavior points to the need to assess Shannon entropy as predictor separately in both types of regions.

In mitochondrial genome, the vast majority of predictors are focused in protein-coding variants with the significant exception of HmtVar disease score, widely used for pathogenic detection in this genome. Looking to benchmark outcomes in mitochondrial SNVs, locus variability as Shannon entropy presented the best performance for the classification of SNVs in non-coding regions. Conversely, the behavior of Shannon entropy in protein-coding variants was considerably poor, due to the genetic code redundancy. This result, joined with the fact that Shannon entropy also presented a low overall performance in not-splice SNVs (AUC=0.86), corroborate the unsuitability of this predictor for SNVs placed in codonic sites. Nevertheless, Splicing process carries an additional restriction for splice-sites that can be easily translated in to locus variability terms, confirmed by its performance as predictor in this positions (AUC=0.95).

Compared with ada score and rf score, locus variability as Shannon entropy matches or even surpassed their outcomes. These two predictors are machine learning ensemble scores, that combine the information from several predictors (Position Weight Matrix (PWM) model, MaxEntScan (MES), Splice Site Prediction by Neural Network (NNSplice), GeneSplicer, Human Splicing Finder (HSF), NetGene2, GENSCAN and SplicePredictor) for SNVs located in splicing consensus regions^21^. Ada score is the result of training an adaptive boosting model while rf score is a similar machine learning approach using a random forest. Both predictors are interesting tools but has the non trivial limitation that there is a limited number of pre-computed values for these scores. In this sense, Shannon entropy is a measurement of locus variability that can be easily calculated from gnomAD variant allele frequencies extracted for any specific genomic position and could be computed at genome-wide scale, presenting a clear advantage over these two scores, with out loosing sensibility, specificity or precision.

As locus variability can be understood as the final state of the evolutionary process, in this study we also compared Shannon entropy against two different conservation scores as phyloP and phastCons, that relies in different strategies. PhastCons is a hidden Markov model-based method that estimates conservation rate, for a specific site, taking in to account the rates of neighboring sites. By contrast, PhyloP scores measure evolutionary conservation at individual alignment sites, giving information not only about the magnitude but also about the direction of the evolution rate compared with a neutral drift model. The two methods have different strengths and weaknesses, PhastCons is effective for conserved elements/regions detection and phyloP, on the other hand, is more appropriate for evaluating signatures of selection at particular nucleotides or classes of nucleotides. Shannon entropy outperformed clearly both conservation scores for pathogenic condition detection. Therefore, this locus variability displays a map among genome-wide splice-sites, that inform about the differential selective pressure for each position, adding granular information that escape to long run evolutionary measurements.

Regarding to benchmark results in mitochondria, it seems that the locus variability in non-coding sequences shows a straight forward relationship with deleteriousness, not observed in coding sequences. Most of the predictors used in mitochondrial SNVs are designed for protein-coding variants, not for variants located at regulatory regions, t-RNA or r-RNA genes. For these specific regions, Shannon entropy is presented as an interesting predictor for pathogenic SNVs, surpassing another predictor widely used as is HmtVar disease score. As observed in splice-sites, Shannon entropy also surpassed conservation scores as pathogenic predictors, in mitochondrial non-coding regions.

To conclude, Shannon entropy can act as an accurate predictor tool that matches or even surpasses pathogenic mutation detection compared with state-of-the-art scores, for SNVs placed in genomic positions that escape the effect of the redundancy of genetic code. In consequence, it could be integrated in variant ranking protocols, in order to reduce the number SNVsclassified as variants of uncertain significance in the context of mendelian or mitochondrial diseases.

## WEB RESOURCES

R statistical software, https://www.r-project.org/

Variant Effect Predictor, https://www.ensembl.org/Tools/VEP/

VariBench, http://structure.bmc.lu.se/VariBench/

